# All families of transposable elements were active in the recent wheat genome evolution and polyploidy had no impact on their activity

**DOI:** 10.1101/2022.11.25.517938

**Authors:** Nathan Papon, Pauline Lasserre-Zuber, Hélène Rimbert, Romain De Oliveira, Etienne Paux, Frédéric Choulet

**Affiliations:** Université Clermont Auvergne, INRAE, GDEC, 63000, Clermont-Ferrand, France; Gencovery, 69000, Lyon, France; VetAgro Sup Campus agronomique, 63370, Lempdes, France

**Keywords:** *Triticeae*, genome evolution, transposable elements, structural variations, polyploidy

## Abstract

Bread wheat (*Triticum aestivum* L.) is a major crop and its genome is one of the largest ever assembled at reference-quality level. It is 15 Gb, hexaploid, with 85% of transposable elements (TEs). Wheat genetic diversity was mainly focused on genes and little is known about the extent of genomic variability affecting TEs, transposition rate, and the impact of polyploidy. Multiple chromosome-scale assemblies are now available for bread wheat and for its tetraploid and diploid wild relatives. In this study, we computed base pair-resolved, gene-anchored, whole genome alignments of A, B, and D lineages at different ploidy levels in order to estimate the variability that affects the TE space. We used assembled genomes of 13 *T. aestivum* cultivars (6x=AABBDD), *T. durum* (4x=AABB), *T. dicoccoides* (4x=AABB), *T. urartu* (2x=AA), and *Aegilops tauschii* (2x=DD). We show that 5 to 34% of the TE fraction is variable, depending on the species divergence. Between 400 and 13,000 novel TE insertions per subgenome were detected. We found lineage-specific insertions for nearly all TE families in di- tetra- and hexaploids. No burst of transposition was observed and polyploidization did not trigger any boost of transposition. This study challenges the prevailing idea of wheat TE dynamics and is more in agreement with an equilibrium model of evolution.

## Introduction

Transposable elements (TEs) are key factors of genome evolution and their contribution to plant phenotypic variations and adaptation were shown in many studies (for reviews:(Lisch, 2013; Baduel and Quadrana, 2021). They are particularly prominent in the genomes of *Triticeae*, a tribe of monocot plants encompassing important crops like wheat, barley, and rye, which diverged from a common ancestor ∼13 MYA. *Triticeae* genomes contain millions of copies of TEs, making them a good model to understand the dynamics of TEs in complex genomes, their contribution to structural variations (SVs) and species adaptation. Bread wheat (*Triticum aestivum*) is the most widely grown crop on earth. Its genome is hexaploid and was shaped by two successive events of allo-polyploidization (reviewed recently in (Levy and Feldman, 2022) that brought together three related diploid subgenomes called A, B, and D, originated from three diploid species which common ancestor was estimated around 6 MYA (Marcussen et al., 2014; Middleton et al., 2014; Glémin et al., 2019; Avni et al., 2022; Li et al., 2022). They share a common karyotype with 7 pairs of chromosomes representing around 5 Gb each. A first interspecific hybridization event occurred ∼0.8 MYA between two *Triticeae* diploid species from the A and B lineages, and a second event occurred between a tetraploid (AABB) and a diploid species carrying a D genome during the period of wheat domestication 9000 YA. The closest living representative diploid species of AA, BB, and DD are *Triticum urartu* (AA), *Aegilops speltoides* (BB), and *Aegilops tauschii* (DD) which divergence time with A, B, D wheat subgenomes were estimated to 1.3, 4.4, and 0.01 MYA, respectively. Wild and cultivated tetraploid wheats (AABB) are *Triticum turgidum* ssp. *dicoccoides* and ssp. *durum*, respectively.

The large size of its genome (∼5 Gb per haploid subgenome), mainly explained by a massive TE content (85%), has limited for decades our capability to assemble them and, thus, characterize their content, organization, and diversity. Before assessing a complete genome sequence, studies showed high variation and sub-genome specificity of TEs (Yaakov et al., 2013a; Yaakov et al., 2013b). Evidence of TE mobilization in newly formed polyploids was described for a few families as well as changes in the epigenetic status (Kraitshtein et al., 2010; Yaakov and Kashkush, 2011, 2012). These studies tended to conclude that polyploidy was a genomic shock and suggest that TEs evolve by bursts where a few families escape to silencing and expand massively in a short time period, leading to rapid genome diversification. In 2018, a reference-quality genome assembly was produced for the hexaploid cultivar Chinese Spring (IWGSC, 2018) and a deep comparative analysis of the ∼4 million TE copies shaping the A, B, and D subgenomes led to unexpected conclusions about the evolutionary dynamics of TEs (Wicker et al., 2018). Since their divergence 6 MYA, there was a near-complete TE turnover between A-B-D, so that ancestral TEs have been lost while more recent ones have amplified independently in the three diploid lineages, and no burst of TE transposition was observed after polyploidization events. Surprisingly, despite the TE turnover, the wide majority of the ∼500 TE families are still present in similar proportions in each subgenome, although they evolved into subgenome-specific variants i.e., subfamilies. Abundant families are still the same while low-copy families persist at low-copy numbers. Hypotheses raised were that TE turnover is highly regulated and may follow evolutionary rules to maintain a functional genome architecture. These conclusions challenged the “burst”-centered view of plant TE dynamics and is rather in line with an alternative scenario of equilibrium as observed, for instance, in *Brachypodium distachyon* natural populations where TE activity is “remarkably constant” (Stritt et al., 2018) or as exemplified by the Alesia family (Stritt et al., 2021) which maintains at low-copy numbers in the Angiosperms across the generations, suggesting self-regulatory mechanisms (Cosby et al., 2019). TEs might maintain at an equilibrium between deletion and amplification in *Triticeae* but it is still a matter of debate (Bariah et al., 2020).

The limit with comparing A-B-D genomes is that all ancestral TEs have been erased so that it is not possible to trace recent deletion/transposition events. For that, it is necessary to compare genomes that have diverged much more recently and identify structural variations. This is what was performed by producing resequencing data on flow-sorted chromosomes 3B in a panel of 45 diverse *Triticeae* (De Oliveira et al., 2020). The extent of TE presence-absence variations (PAVs) was estimated to 7-8% per *T. aestivum* accession and up to 24% in more distant species and, again, no burst of any specific TE family was observed. In contrast, recent TE insertions were detected for a wide diversity of families, confirming that most of families were active and transposed. Recently, several *Triticeae* genomes have been assembled at reference-quality level, offering the opportunity to analyze TE dynamics at a resolution never reached so far using whole genome alignments. Analysis of five full-length LTR-retrotransposon families across *T. aestivum* assembled genomes confirmed that distinct subfamilies were active in diploids mainly, in waves lasting hundred thousand years, with only sparse evidence of recent insertions in polyploids (Wicker et al., 2022). However, the extent of variability due to TEs deletion/amplification is still unknown.

In this study, we developed a method in order to compute *Triticeae* whole-genome sequence alignments guided by anchor-genes and retrieve the complete landscape of TE variability originated from deletions and transpositions. We thus compared the fully assembled sequences of the A genomes, the B genomes, and also the D genomes between di- tetra- and hexaploids: *T. urartu* (AA; (Ling et al., 2013)), *Ae. tauschii* (DD; (Jia et al., 2013)), *T. dicoccoides* (AABB; (Avni et al., 2017)), *T. durum* (AABB; (Maccaferri et al., 2019)), and 13 *T. aestivum* (AABBDD; (IWGSC, 2018; Walkowiak et al., 2020; Aury et al., 2022)). We show that variability affects from 5 to 34% of the TEs depending on the species compared. We identified 51,928 recent transposition events involving 346 different families that were active recently in all species whatever their ploidy level. We show that transposition rate is similar in di- tetra- and hexaploids, confirming the equilibrium we observed previously, and confirming that polyploidy did not disturb this equilibrium.

## Material and Methods

### Genome sequence data

The reference-quality assembled genome sequences of *T. urartu* cultivar G1812 v2.0 (PRJNA337888) (Ling et al., 2018), *Ae. tauschii subsp. strangulata* cultivar AL8/78 v4.0 (PRJNA182898) (Luo et al., 2017), *T. turgidum subsp. durum* cultivar Svevo v1.0 (PRJEB22687) (Maccaferri et al., 2019), *T. turgidum subsp. dicoccoides* isolate Atlit2015 ecotype Zavitan v2.0 (PRJNA310175) (Zhu et al., 2019) and *T. aestivum* RefSeq v1.0 (PRJEB27788) (IWGSC, 2018) were downloaded from NCBI (https://www.ncbi.nlm.nih.gov/). Reference-quality assembled genome sequences of *T. aestivum* accessions ArinaLrFor, CDC Landmark, CDC Stanley, Jagger, Julius, LongReach Lancer, Mace, Norin61, Spelt, and SY_Mattis were downloaded from https://wheat.ipk-gatersleben.de/ (Walkowiak et al., 2020). We also used the Renan genome sequence that we published previously (GCA_937894285) (Aury et al., 2022), and that of Tibetan wheat Zang1817 (Guo et al., 2020).

### TE annotation and comparison of family proportions

We used the available TE annotations that we computed previously for Chinese Spring (Wicker et al., 2018; Aury et al., 2022) and Renan (Wicker et al., 2018; Aury et al., 2022). For all the other genome sequences compared in this study, we did not use the available annotation but rather produced a TE annotation using CLARITE and the ClariTeRep library (Daron et al., 2014) with the same parameters as for Chinese Spring and Renan. CLARITE uses RepeatMasker (Smit et al., 1996-2004) for similarity search, applies a step of defragmentation in order to merge adjacent predictions that describe a single element, and eventually applies a step of reconstruction of nested insertions. The abundance of each TE (sub)family in a genome was calculated by cumulating the length of all fragments assigned a given (sub)family divided by the total length of TEs. To investigate differences of TE family abundance between genomes, and potential enrichment in the variable fraction of the genome, we calculated proportions of each TE family (cumulated length assigned to a given family divided by the subgenome size) and computed log2 ratios. Only families accounting for more than 100 kb in the analyzed subgenome were considered.

### Estimation of the extent of genomic variability using ISBP markers

Based on CLARITE predictions of TEs, we extracted 150 bps encompassing the 5’ and 3’ junctions of each TE extremity with its insertion site with 75 bps on each side of the junction as previously described (De Oliveira et al., 2020). These 150 bps tags correspond to ISBP markers (Insertion Site Based Polymorphism (Paux et al., 2006)) that are expected to be unique kmers at the whole genome level. For each genome studied, we extracted all ISBPs and discarded those containing Ns. In case two ISBPs overlap by 100 bps or more, we kept only one of both. We also discarded ISBPs that are non-unique in the genome from which they were designed: mapped with Minimap2 (Li, 2018) at multiple loci with cutoff 98% identity and 100% query overlap. Presence/absence of ISBPs were searched in all compared genomes using Minimap2 option -xsr -w5. An ISBP was considered absent (PAV) if no match was found considering at least 95% identity and 95% coverage (maximum 7 bases soft-clipped). Only pseudomolecule sequences were considered for this analysis, meaning that unanchored scaffolds (“ChrUn”) were excluded. To determine nucleotide positions defining the limits between distal (R1 and R3), proximal (R2), and (peri)-centromeric (C) regions of each chromosome (as previously defined in Chinese Spring (IWGSC, 2018)) in all species/accessions analyzed, we used the Chinese Spring ISBP mapping data. Hence, the chromosomal position of the ISBP that was the closest to a border (between R1 and R2 of chr1A for instance) defined in CS was used as border in the compared species/accessions.

### Identification and selection of orthologous intergenic regions

We used the 105,200 High Confidence (HC) genes with a position along the 21 pseudomolecules of RefSeq v1.1 (IWGSC, 2018) as anchors to find orthologous intergenic regions in other genomes. The nucleotide sequences of the corresponding CDSs were mapped using GMAP (v18.05.11, (Wu and Watanabe, 2005), options: gmapl –cross_species) on the homeologous chromosome for each genome compared. Only best hit with at least 90% identity and 90% coverage were considered and kept for the subsequent analyses. We developed a Python script (Get_Collinear_Region.py) in order to retrieve all orthologous intergenic regions (oIGRs) that were flanked by the same pair of neighbor orthologs (collinearity) in Chinese Spring and in the compared genome. Cases of tandem duplicated gene copies that mapped at the same positions in a compared genome were dealt with by the script so that it specifically determined which copies delimitate an oIGR. Chromosome positions of these pairs of neighbor orthologs were used to extract the corresponding genomic segments (including the genes at both extremities of oIGRs) in both compared genomes. oIGRs were then aligned with BLASTN (v2.11.0+, (Camacho et al., 2009), threshold evalue: 1e-5) and we filtered out HSPs (High Scoring Pairs) with a cutoff at 90% identity. We excluded HSPs which coordinates were included into a larger one. This happened in case of lineage-specific tandem duplications, or when a novel copy of an element shared homology with another TE present within the aligned region. Coordinates of HSPs were then used to create BED files and we used “Bedtools merge” (bedtools/2.26, (Quinlan and Hall, 2010)) to merge overlapping conserved segments for each genome. We then used “Bedtools complement” to create BED files describing the variable (i.e., specific) segments between genomes i.e., sequences that are subject to presence-absence variations (PAVs) between two compared oIGRs. PAV candidates corresponding to gaps in the assembly (stretches of Ns) were identified and discarded from PAVs. We eventually used “Bedtools intersect” to retrieve TE annotation of the conserved/variable sequences.

### Detection of recent TE insertions and estimation of the insertion dates

BED files describing the positions of PAVs (see above) were analyzed in the objective of finding PAV coordinates that fit nearly perfectly (i.e., with a tolerance of 10 bps at 5’ and 3’ extremities) with the coordinates of a TE, with status “complete” annotated by CLARITE, as evidence for recent transposition (or possible excision for class 2 elements). For each candidate of novel insertion, we searched for the presence of a target site duplication (TSD) as molecular evidence of insertion. TSD are short motifs repeated at 5’ and 3’ ends of a TE and immediately flanking the terminal motifs that determine the exact borders of the TE. Since the predicted extremities of TEs may not correspond exactly to the terminal nucleotides, TSDs were searched within a subsequence of 20 nucleotides (nts) overlapping both TE extremities: 10 nts on each side of the predicted extremity for superfamilies RLG, RLC, RLX, and DTC; 5 nts inside+10 nts outside for superfamilies DTM, DTT, DTH, and DTA. We developed Python scripts (1 per superfamily; available at https://forgemia.inra.fr/umr-gdec/scripts_files/) to identify TSDs of variable size, depending on the superfamily considered (Wicker et al., 2007), immediately flanking terminal motifs: 5’-TG and CA-3’ for RLGs, RLCs, and RLXs; 5’-CACT[AG] and [CT]AGTG-3’ for DTCs. For the other superfamilies with undefined terminal motifs, the entire subsequence of 20 nts was scanned for the presence of a TSD. The expected TSD size was 5 nts for RLGs, RLCs, and RLXs, 3 nts for DTCs and DTHs, 8 nts for DTAs, varying between 9 and 11 nts for DTM, and a TA duplication was expected for DTTs. We tolerated one SNP between 5’ and 3’ TSDs. No TSD was searched for LINEs/SINEs. Insertion dates of LTR retrotransposons were estimated by aligning the two LTRs of an element with BLASTN and we used a mutation rate of 1.3×10^−8^ substitutions/site/year (SanMiguel et al., 1998) as previously described (Wicker et al., 2018). Distances of newly inserted TE from the nearest predicted gene were computed with “Bedtools closest”.

## Results

### TE annotation and comparison of A, B, and D subgenomes between di, tetra- and hexaploid *Triticeae*

To avoid biases due to different TE annotation approaches, we predicted TEs in the genome sequences of *T. aestivum* (13 accessions), *T. urartu, Ae. tauschii, T. dicoccoides* and *T. durum* (abbreviated *Tae, Tur, Aet, Tdi, Tdu*, respectively) with the same method and criteria, using CLARITE (Daron et al., 2014). TEs represent between 86 and 87% for the A lineage, between 84 and 85% for B, and between 82 and 83% for D. We previously showed that A-B-D diverged ∼5-6 MYA, a time during which the TE turnover has erased ancestral TEs so that there is (almost) no TE conserved between orthologous loci (homeologous in polyploids). Here, we focused our work on a much more recent evolutionary scale, by comparing A subgenomes together, B together, and D together, at different ploidy levels. TEs shaping the intergenic space are, thus, conserved because they were present in the common ancestor of the genomes compared.

The amount of TEs appeared very similar, even of each TE superfamily, when we compared A (sub)genomes, so did B, and D (Table 1; Supplemental Table S1). *T. urartu*^*A*^ appeared the most different, which fits with its earlier divergence (1.3 MYA). To get a first flavor about the extent of variability, we started by comparing (sub)genome-wide proportions of TE families between lineages. Using Chinese Spring (CS) as a reference, we distinguished 301 TE families (those accounting for more than 100 kb in at least 1 subgenome), with the 20 most abundant representing >84% of the TE fraction, meaning that most families are present at low-copy number. We showed that 97% (292/301) of the families were present in similar proportions in the compared genomes (fold-change<2), suggesting that none of these *Triticeae* have experienced a massive transposition “burst” of any family since they diverged, neither before, nor after, the polyploidization events. Rare cases of differential abundance were observed for nine very low-copy families (DTM_famc13, RIX_famc25 XXX_famc10-150-46-53-57-60, DHH_famc1) which abundance varies around the 100 kb cutoff applied here and for which the extent of the difference may not be associated with TE amplification but rather to methodological limits (partial genome assembly and anchoring, TE annotation). The only remarkable case is family XXX_famc10 which is not a TE *per se* but rather a subtelomeric satellite repeat specific to the B lineage. It exhibited strong differences of copy number affecting the A subgenome between diploid *Tur*^*A*^ and tetraploid *Tdi*^*A*^ which is explained by the previously characterized 4A/7B translocation (Dvorak et al., 2018) that brought a part of the B genome onto the 4A chromosome in the tetraploid ancestor. Such global conservation of family proportions may hide higher levels of structural variations between those genomes which we, thus, investigated using uniquely mappable TE-derived markers.

**Table 1:** Proportions of TE superfamilies in the A, B, and D subgenomes of *Triticeae* (sub)genomes annotated with CLARITE. Proportions are expressed as the percentage of sequences assigned to each superfamily relatively to the (sub)genome size. TIRs: terminal inverted repeats.

|  | *T.*<br>*urartu* | *T. dicoccoides* |  | *T. durum* |  | *Ae.*<br>*tauschii* | *T. aestivum*<br>(Chinese Spring) |  |  |
| --- | --- | --- | --- | --- | --- | --- | --- | --- | --- |
| (sub)genome | A | A | B | A | B | D | A | B | D |
| All TEs | 86.8% | 85.4% | 84.4% | 86.9% | 86.0% | 83.4% | 86.1% | 84.7% | 83.1% |
| Class I: Retrotransposons | 71.8% | 70.7% | 66.5% | 72.0% | 67.9% | 62.0% | 71.9% | 67.4% | 62.4% |
| RLG (Gypsy) | 49.7% | 49.7% | 46.0% | 50.8% | 47.0% | 40.7% | 50.9% | 46.8% | 41.4% |
| RLC (Copia) | 18.6% | 17.3% | 16.0% | 17.6% | 16.3% | 16.4% | 17.5% | 16.2% | 16.3% |
| RLX (unclassified) | 2.5% | 2.7% | 3.4% | 2.7% | 3.5% | 3.7% | 2.6% | 3.5% | 3.7% |
| RIX (LINE) | 0.94% | 0.95% | 1.10% | 0.95% | 1.13% | 1.08% | 0.82% | 0.96% | 0.93% |
| RSX (SINE) | 0.01% | 0.01% | 0.01% | 0.01% | 0.01% | 0.01% | 0.01% | 0.01% | 0.01% |
| Class II: DNA transposons | 14.4% | 14.2% | 17.0% | 14.3% | 17.2% | 20.7% | 13.7% | 16.5% | 20.1% |
| DTC (CACTA) | 13.4% | 13.2% | 15.9% | 13.3% | 16.1% | 19.4% | 12.8% | 15.5% | 19.0% |
| DTA (hAT) | 0.01% | 0.01% | 0.01% | 0.01% | 0.01% | 0.01% | 0.01% | 0.01% | 0.01% |
| DTM (Mutator) | 0.35% | 0.34% | 0.43% | 0.35% | 0.44% | 0.55% | 0.30% | 0.38% | 0.48% |
| DTT (Mariner) | 0.15% | 0.15% | 0.17% | 0.15% | 0.18% | 0.19% | 0.14% | 0.16% | 0.17% |
| DTH (Harbinger) | 0.17% | 0.17% | 0.17% | 0.17% | 0.18% | 0.19% | 0.15% | 0.16% | 0.18% |
| DTX (unclassified with TIRs) | 0.23% | 0.23% | 0.23% | 0.23% | 0.23% | 0.24% | 0.21% | 0.20% | 0.22% |
| DHH (Helitron) | 0.05% | 0.05% | 0.08% | 0.05% | 0.08% | 0.06% | 0.05% | 0.08% | 0.05% |
| DXX (unclassified class II) | 0.01% | 0.01% | 0.01% | 0.01% | 0.01% | 0.01% | 0.00% | 0.00% | 0.00% |
| Unclassified repeats XXX | 0.59% | 0.61% | 0.89% | 0.60% | 0.85% | 0.74% | 0.51% | 0.81% | 0.59% |
| Genes and unannotated DNA | 13.2% | 14.6% | 15.6% | 13.1% | 14.0% | 16.6% | 13.9% | 15.3% | 16.9% |

### Extent of structural variations affecting TEs through the mapping of ISBP markers

Although TEs are repeated, each copy is inserted into a different locus, which makes the 5’ and 3’ junctions between TE extremities and the insertion site unique kmers at the whole genome level. We used this valuable feature to address presence/absence variations (PAVs), also called TIPs (transposon insertion polymorphisms) or ISBPs (Insertion Site-Based Polymorphisms) among the wheat research community, between orthologous loci. We extracted 150 bps encompassing each TE extremity of all genomes which provided a dataset of, on average, 1.7, 1.9, and 1,5 million uniquely mappable ISBP markers from the A, B, and D subgenomes, respectively. ISBPs mapped with at least 95% identity over 95% of their length were considered present while no match revealed the absence i.e., PAV. Proportions of markers subject to PAVs in all pairwise comparisons are presented in Figure 1.

**Figure 1:**
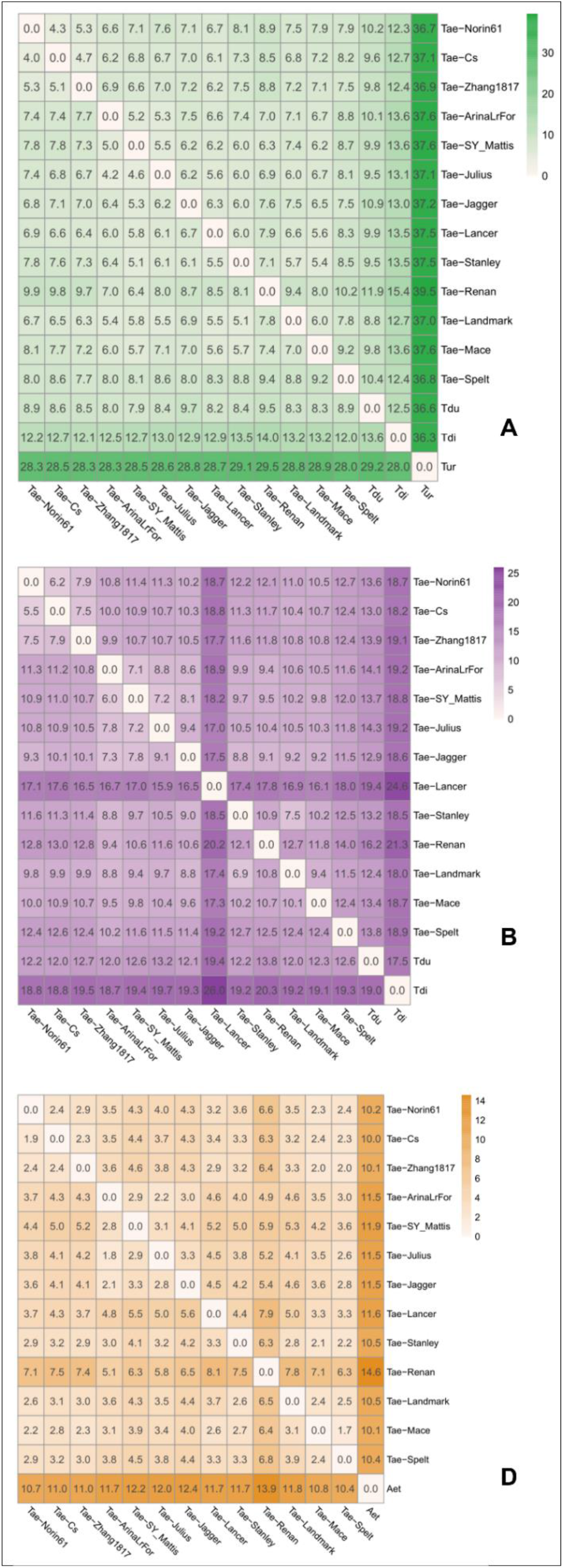
Matrix of the levels of variability (PAVs) estimated using ISBP mapping. Values represent the percentages of ISBP markers that are under PAVs in pairwise comparisons i.e., present in the query (in rows) and absent in the reference (in columns) genome.

At the interspecific level, considering CS as a reference for *T. aestivum*, we show that TE PAVs ranged from 10% for the D subgenome (*Tae*^*D*^ *vs Aet*^*D*^) to 37% for the A subgenome (*Tae*^*A*^*-Tdi*^*A*^*-Tdu*^*A*^ *vs Tur*^*A*^), reflecting the earlier divergence of *T. urartu*. Comparing tetraploids to hexaploids showed that *T. aestivum* is closer to *T. durum* than to *T. dicoccoides*. In addition, the variability is higher for the B than for A subgenome: proportions were estimated to 10% (A) and 13% (B) for *Tae* compared to *Tdu*, and to 13% (A) and 18% (B) compared to *Tdi*. We observed the same shift between A and B genome variability when comparing tetraploids together (*Tdi versus Tdu*). Altogether, these data highlighted that around 10-18% of the TE space is variable (specific), in terms of presence/absence, between species *T. aestivum, T. durum, T. dicoccoides*, and *Ae. tauschii*, while it is much variable compared to *T. urartu*.

At the intraspecific level, ISBP-based PAV detection using 13 fully assembled genomes of *T. aestivum* revealed a similar pattern: D subgenome is the least variable (2-5%), A is intermediate (4-10%), and B is the more variable (6-13%). Outliers were observed for the B subgenome of CDC Lancer (16%) due to the presence of an alien chr2B originating from *Triticum timopheevii* (Walkowiak et al., 2020). A slight increase (1-2%) of variability was observed for Renan compared to all other European wheat accessions, due to the sequencing method that was different (Oxford Nanopore *versus* Illumina) and may have led to 1-2% ISBPs with sequencing errors (indels preventing from aligning 95% of the ISBP length).

PAV levels detected here were in agreement with the existence of two phylogenetically distinct groups corresponding to the Asian (CS, Norin61, Zang1817) and European wheat genetic pools. Indeed, variability was slightly lower within each group (A: 5-6%; B: 7-9%; D: 2-4%) than between groups (A: 7%; B: 11%; D: 4%).

These global estimates at the whole subgenome level may hide strong local differences. Chromosome extremities (distal regions) were, on average, four times more variable than the central regions of chromosomes (borders as defined in (IWGSC, 2018)). For instance, between the closely related Asian accessions CS and Norin61 (Fig. 1), 4% of TEs are variable, however, it reached up to 9% in the fast-evolving distal regions whereas it is limited to 2% for the rest of the genome. Hence, TE PAVs can be high (>10%) in the distal regions even between accessions that are closely related. These results confirmed, at a short evolutionary scale, previous assessments suggesting accelerated evolution in the recombinogenic distal regions compared to the rest of the genome.

This ISBP-based PAV detection method provided an unbiased genome-wide view of the extent of the variable *versus* conserved parts of *Triticum*/*Aegilops*. Roughly, it represents 5-10% and 10-20% of the TE-derived markers at the intra- and inter-specific levels, respectively, in pairwise comparisons, with substantial differences between the B (more variable), A (intermediate), and D (least variable) subgenomes. After getting this first flavor of genome-wide structural variability using ISBPs as proxy, we established a method to characterize at a bp resolution which TEs comprise the variable and conserved parts through pairwise alignments of orthologous intergenic regions.

### TE variability assessed by whole genome alignments

Aligning Gb-sized genomes containing 85% of TEs is not trivial. In this regard, we developed a dedicated approach aiming at identifying collinear orthologous genes in order to target the pairwise alignment of pre-identified orthologous intergenic regions (oIGRs). Therefore, we mapped the 105,200 predicted genes of *T. aestivum* Chinese Spring (35,345, 35,643 and 34,212 carried by subgenomes A, B and D, respectively) on related genomes with high stringency and found unambiguous orthologs for 79%, 96%, 94%, and 94% of them in *T. urartu*^*A*^, *Ae. tauschii*^*D*^, *T. dicoccoides*^*AB*^, and *T. durum*^*AB*^, respectively. These genes were used as anchors to guide all genome sequence alignments. From this, we extracted all intergenic regions flanked by collinear neighbor orthologs, representing a dataset of 18,028, 30,688, 59,462, and 60,230 oIGRs, respectively. Their cumulated length represents 34% (1.6 Gb) of the *Tur*^*A*^/*Tae*^*A*^ subgenomes, 95% (3.6 Gb) of the *Aet*^*D*^*/Tae*^*D*^ subgenomes, 90% (4.3 Gb) of the *Tdi*^*AB*^*/Tae*^*AB*^ genomes, and 91% (4.3 Gb) of the *Tdu*^*AB*^*/Tae*^*AB*^ genomes. We then retrieved the positions of the conserved *versus* variable (specific) sequences from pairwise oIGR alignments as illustrated in Figure 2. This fine-tuned approach allowed us to get a bp-resolved view of TE copies that are conserved or affected by PAVs (due to insertions, duplications, or deletions) among >90% of the A-B-D subgenomes, except with *Tur*^*A*^ which exhibited lower collinearity and lower assembly quality.

**Figure 2:**
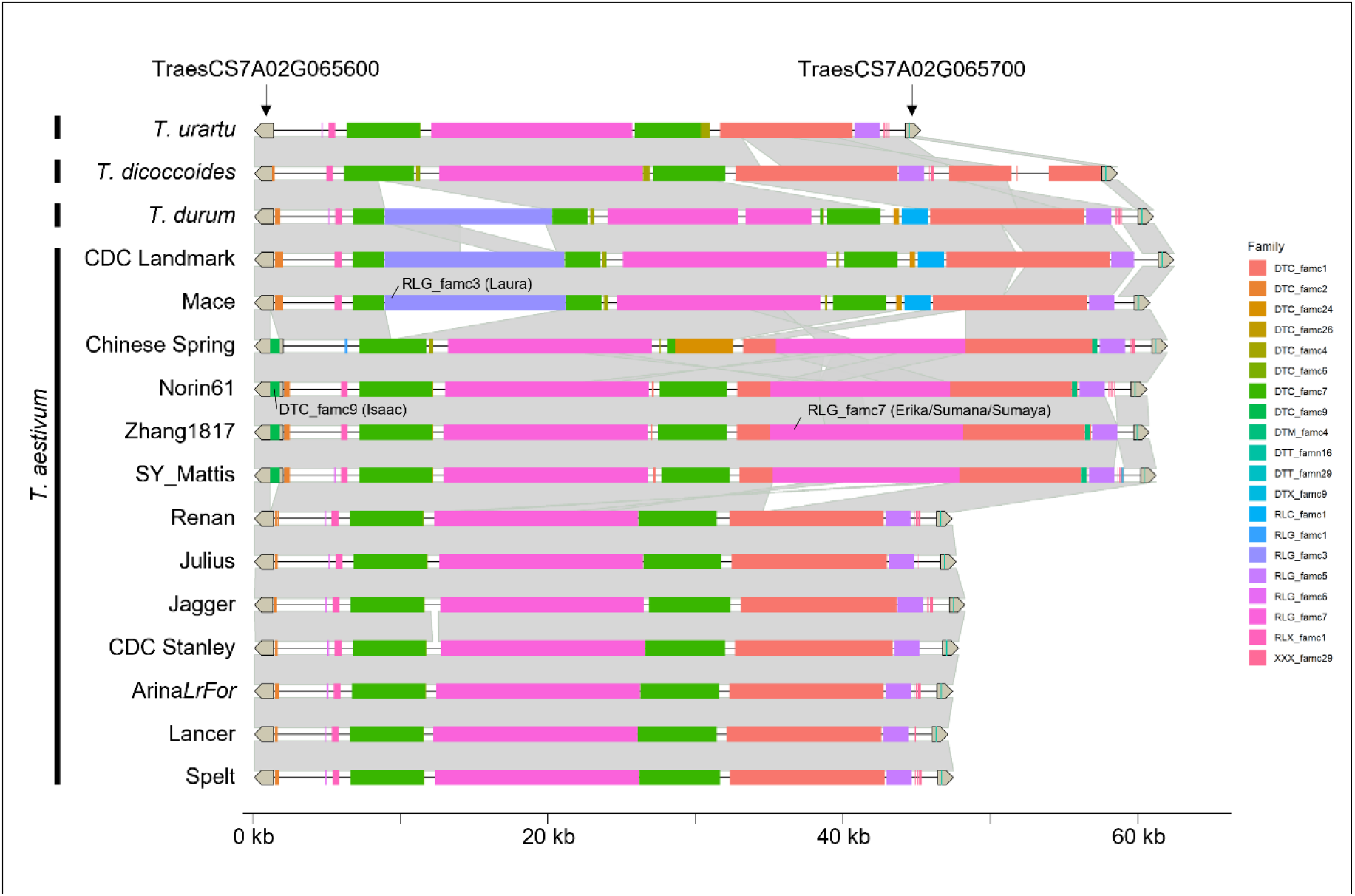
Example of an orthologous intergenic region on chr7A compared using Minimap2 across the four species with (sub)genome A (*T. urartu, T. dicoccoides, T. durum, T. aestivum*) and 13 *T. aestivum* accessions. Conserved TEs are covered by grey areas which illuminates regions subject to PAVs. PAV coordinates may fit with TE coordinates suggesting recent TE insertion/excision.

The variable fraction of the A genome represents 34% of CS *Tae*^*A*^ subgenome when compared to diploid *Tur*^*A*^ (Table 2). Tetraploids are much more closely related: the variable fraction represents 7% and 10% of the A subgenome compared to *Tdu*^*A*^ and *Tdi*^*A*^, respectively. The B subgenome exhibits higher variability level: 9% and 13%, respectively. Compared to diploid *Aet*^*D*^, 9% of the *Tae*^*D*^ subgenome is under PAVs. These values are in agreement with the ISBP-based estimates described in the above paragraph although oIGR analysis tended to slightly underestimate the level of variability because we did not sample regions where genes are noncollinear. Considering that A, B, and D subgenomes are shaped by 1.2 million TEs on average, our results revealed that roughly 120k TEs (∼10%) are non-conserved while ∼90% of the TE space is still conserved in pairwise comparisons.

**Table 2:**
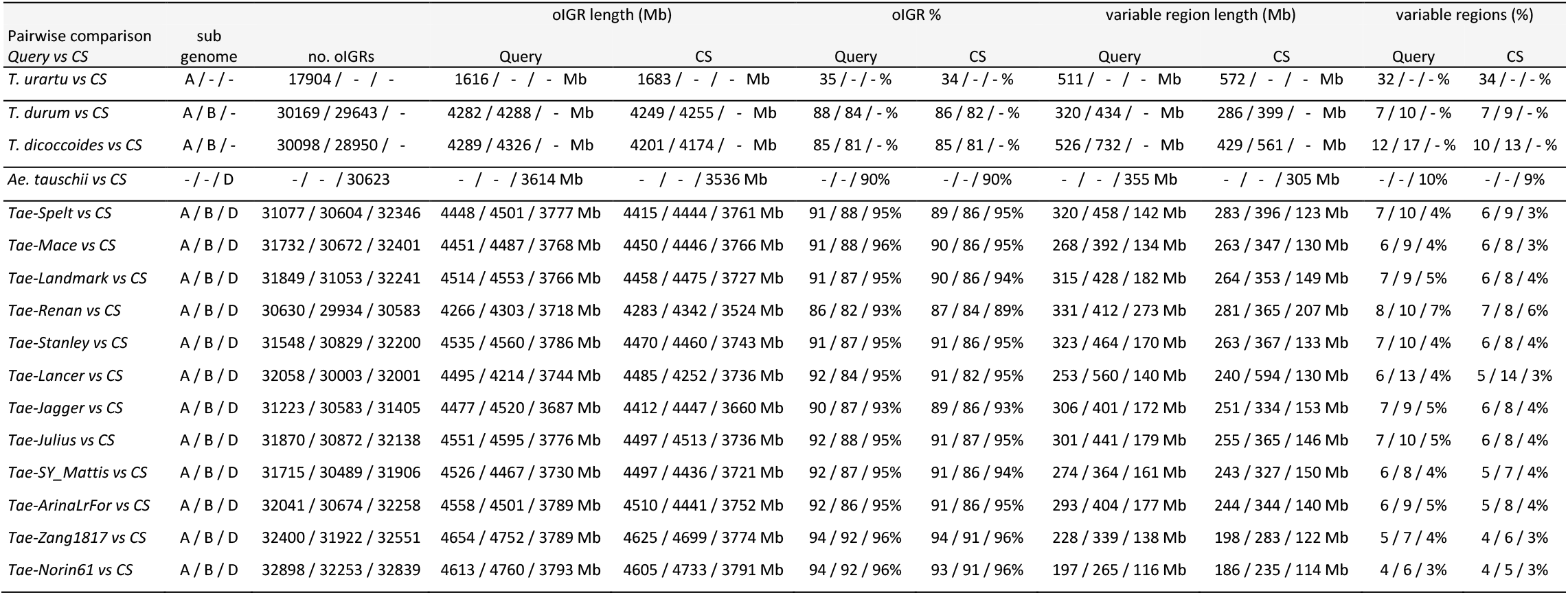
Proportions of the variable TE fraction identified by oIGR alignments in each pairwise comparison with Chinese Spring (CS). *Tae: T. aestivum*.

Intraspecific pairwise comparisons of 13 *T. aestivum* accessions revealed that the variable TE fraction is in the same range than compared to *T. durum*, representing on average 6% of A and 8% of B subgenomes, although it is lower, 4% and 5%, when comparing CS with closer Asian accessions Norin61 and Zang1817. Thus, we did not observe a strong difference in terms of TE turnover when comparing CS with hexaploids and with *T. durum. T. dicoccoides* appeared more distantly related. For the D subgenome, only 4% of CS^D^ TEs were affected by PAVs compared to other accessions which is half that observed compared to *Ae. tauschii* (9%).

Alignments of oIGRs of CDC Lancer revealed a higher level of TE PAVs for the B subgenome due to the presence of an alien chromosome 2B, as commented above. Except for such introgressions, TE variability appeared quite stable and in agreement with SNP-estimated divergence. We did not observe cases where TE activity would have triggered accelerated TE turnover.

Another conclusion we could draw from Table 2 is that the proportions of variable regions are similar in both the query and reference aligned genomes: for instance, 320 Mb (7.5%) of *Tdu*^*A*^ TEs are absent in CS^A^ while, reversely, 286 Mb (6.7%) of the CS^A^ TEs are absent in *Tdu*^*A*^. Between A subgenomes of accessions CS and Norin61, specific TEs account for 186 Mb and 197 Mb, respectively. This shows that none of the genomes analyzed here evolved towards expansion or contraction of the TE space. In contrast, the rate of TE turnover is globally conserved in all lineages analyzed, suggesting that insertions of novel TEs compensate TE loss by deletions.

We then wondered about the composition of the variable fraction of the TE space in order to investigate which families have impacted the recent *Triticeae* genome evolution. Indeed, recent TE amplification of active families could be detected because they are enriched in the variable fraction. Thus, we searched for families whose abundance is substantially enriched in the variable fraction of the genome compared to its genome average. We calculated enrichment ratios for the 100, 113, and 98 most abundant families of the A, B, and D subgenomes, respectively (those representing at least 1 Mb per subgenome in CS). Together they represent 99% of the TE content, the others being low-copy TEs. Fold-changes are shown as heatmaps in Figure 3. The main result here is that such enrichment is rare. The composition of the variable fraction is quite similar to the genome average. However, we found 3 TE families (2 CACTAs and 1 Gypsy) and 2 satellite repeats that were enriched (log2 fold-change>=2.0) in the variable TE fraction of some genomes: RLG_famc8 (Cereba, Quinta), DTC_famc4 (Clifford/Mandrake/Byron), DTC_famc9 (Isaac), and satellites XXX_famc1 (TaiI) and XXX_famc10 (unnamed). RLG_famc8 (Cereba/Quinta) is a centromeric gypsy family which represents on average 1% of the oIGRs but accounts for 5% of the variable sequences detected. Centromeric TEs and telomeric satellites have been, thus, main components of the recent (intraspecific) genome structure diversification. But if these examples are striking, together they were responsible for at maximum 5-6% (in bps) of the PAVs, showing that the observed variability does not originate from amplification bursts of a few very active families. In contrast, the composition of the recently shaped TE space resembles the ancestral one.

**Figure 3:**
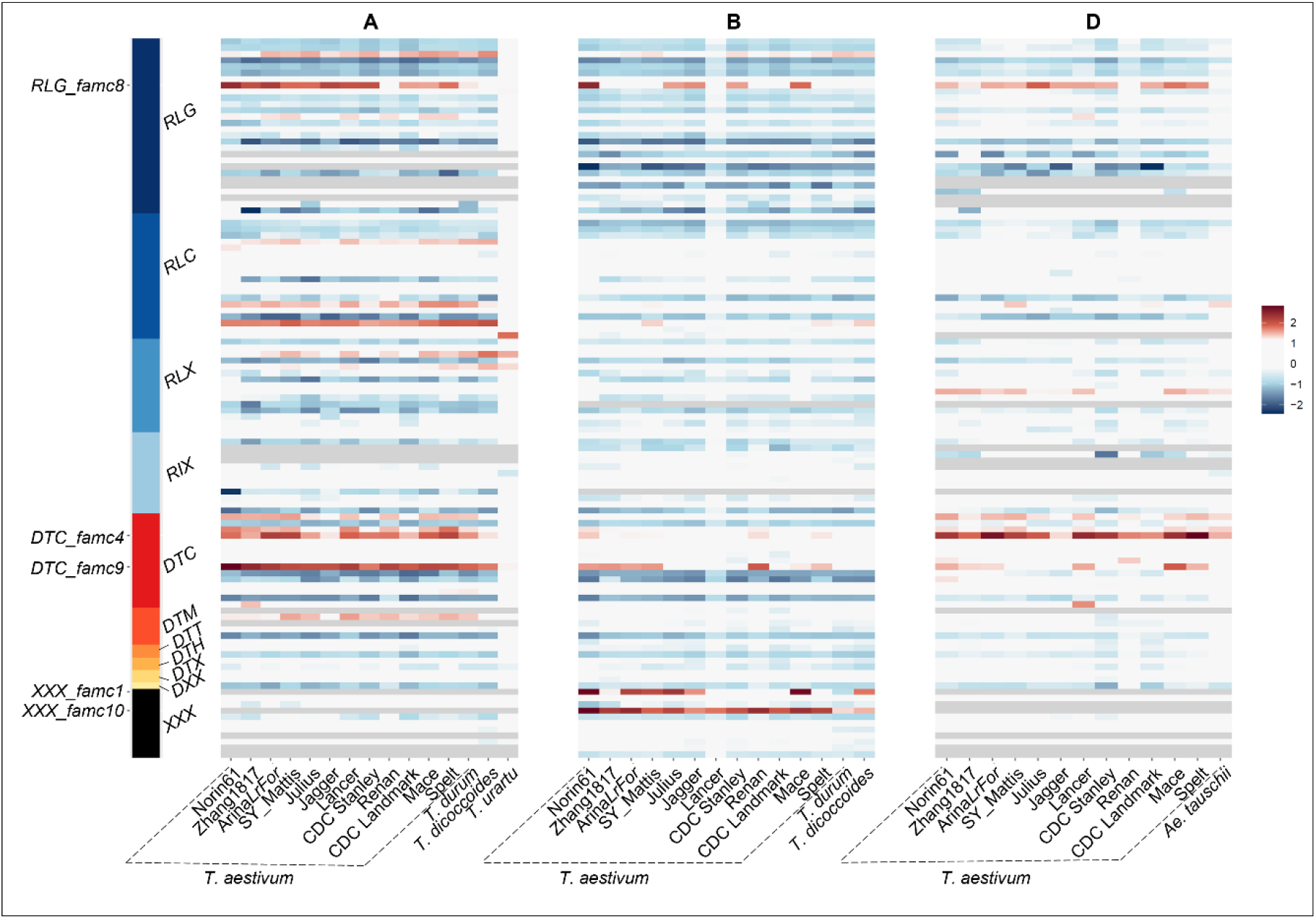
TE family enrichment in the variable fraction of the genome represented as heatmaps for the subgenomes A, B, and D. Enrichment ratios were calculated for the 100, 113, and 98 most abundant families of the A, B, and D subgenomes, respectively (representing at least 1 Mb per subgenome in CS). Abundance of each family (in bps) was retrieved at the whole genome scale and compared to that in the variable fraction of the genome (identified from pairwise comparisons with the reference cultivar Chinese Spring). Log2 ratios between these two proportions were then calculated and represented as heatmaps with red showing families that could be considered as enriched in the variable fraction compared to their genome average. Families were ordered in rows according to their superfamily classification represented by a color code on the left panel and labelled with a 3-letter code. Names of 5 families showing log2 ratios >=2 are indicated on the left.

### Traces of recent transposition events confirms the equilibrium model of TE evolution

Variable regions originated from both deletions and insertions. To get deeper insights into the dynamics of transposition, we searched for PAVs (identified by oIGR pairwise BLAST alignments) which borders fit with borders of a TE, as potential traces of transposition (or potentially excision for class 2 elements). Number of such events are summarized in Table 3. Pairwise comparisons with CS revealed between 400 and 13,000 recent TE insertions per subgenome, depending on the species/accession considered. We found presence of TSDs (target site duplications, i.e., a molecular evidence of transposition) for 78% of them. Density of newly transposed elements follows species divergence time: 6 insertions per Mb in CS^A^ compared to *T. urartu*^*A*^; 2.6 and 3.0 insertions/Mb compared to *T. dicoccoides* A and B subgenomes, respectively; 1.3 and 1.7 insertions/Mb compared to *T. durum* A and B, respectively; 1.3 insertions/Mb compared to *Ae. tauschii*^*D*^. These novel insertions accounted 10-15% of the size of the specific regions identified in each subgenome. At the intraspecific level, we found that A, B, and D subgenomes of CS carry on average 4261, 5460, and 659 specific insertions compared to the 12 other accessions, representing a density of 1.0, 1.2, 0.2/Mb, respectively. Novel insertions were more frequent in the distal regions (as defined in (IWGSC, 2018)) than in the central part of chromosomes e.g., 8.5 *versus* 5.4 novel insertions/Mb in CS^A^ compared to *T. urartu*^*A*^; 1.9/2.0/0.5 *versus* 0.7/1.0/0.1 novel insertions/Mb in CS A/B/D subgenomes compared to other *T. aestivum*.

**Table 3:**
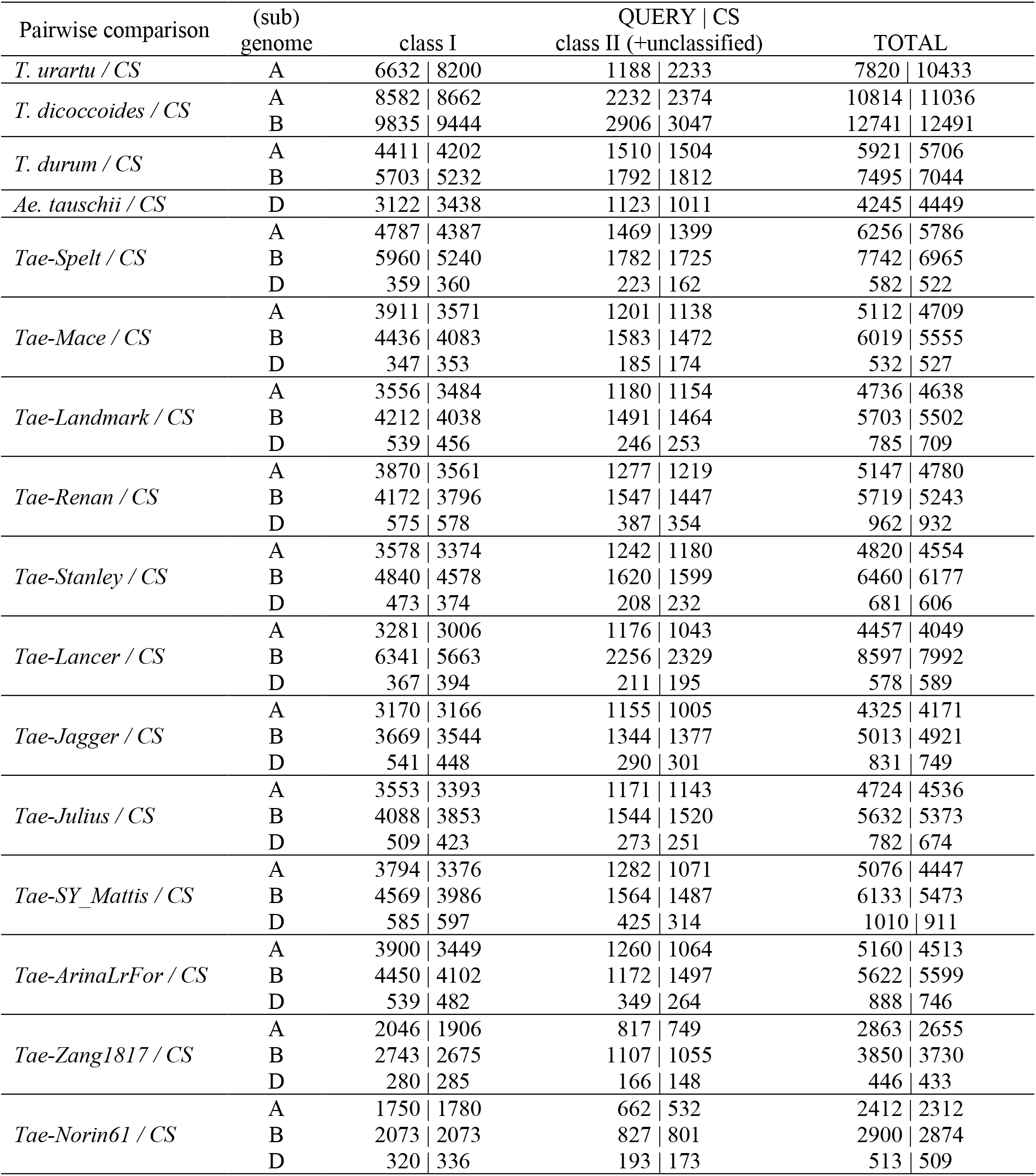
Number of specific TE insertions detected in pairwise oIGR alignments. Number of TEs inserted in the query | subject genomes are indicated relatively to each other.

A striking result is that the numbers of newly inserted TEs were quite similar between the two genomes aligned, whatever the comparison considered (Table 3). For instance, 5706 novel insertions were found in CS compared to the orthologous locus in *T. durum* and, reversely, we detected 5921 novel insertions in *T. durum* compared to CS. Similarly, comparing CS^D^ against *Ae. tauschii*^*D*^ revealed 4245 and 4449 specific insertions, respectively. This was the case for every species/accession compared to the reference. This shows that, firstly, transposition is not silenced in any species analyzed here, with thousands of recent events discovered and, secondly, that transposition rate is somehow constant since the divergence of these genomes. We found no occurrence of enhanced TE amplification in any lineage explored here and found no impact of ploidy on TE activity. In other terms, there were no more transpositions in polyploids than in diploids.

We then wondered which TE families were the most active in this short evolutionary time frame. We found traces of recent transposition for 346 different families, most (79%) of them having transposed in A, B, and D subgenomes. Together these active families represent 99.7% of the whole genome TE content because, although 505 families were distinguished in ClariTeRep, many families were only poorly characterized, with truncated elements, spurious predictions, or misclassified repeats, and we cannot find newly inserted copies for such families. Thus, we applied a 100 kb threshold per subgenome analyzed in order to estimate the proportion of active families in wheat. This retained 301 high confidence families and 89% of them were active. We conclude that virtually all families were active recently and gave rise to newly inserted copies in the recent *Triticeae* evolution. Even at the intraspecific level, we found traces of transposition for 328 families. This situation cannot be explained by cycles of silencing/bursts but is rather in favor of an equilibrium model of evolution.

To go further, we wondered if the level of activity of a given family was different between species/accessions and if the rate of transposition was correlated or not with the abundance of the family. Figure 4 represents the number of specific insertions per pair of aligned subgenomes for the 20 most abundant families. RLC_famc1 (*Angela-WIS*) was the most active family representing around 25% of the recent insertions discovered in all A, B, and D genome comparisons and it is, actually, the most abundant family. But other abundant families DTC_famc2 (*Jorge*), RLG_famc2 (*Sabrina/Derami/Egug*), and RLG_famc1 (*Fatima*) were much less active, representing only ∼1% of the recent insertions. In contrast, RLG_famc3 (*Laura*) and RLG_famc13 (*Latidu*) were among the most active families (reaching 22% of the insertions) while they are less abundant. We conclude that there is no positive nor negative correlation between the recent transposition rate and the family abundance. Again, a striking result is that this pattern is conserved in the compared genomes. For instance, since the divergence of *T. urartu*^A^ and CS^A^, 156 copies of RLG_famc1 (*Fatima*) transposed specifically in *T. urartu*^A^ and 145 in CS^A^. Comparing *Ae. tauschii*^*D*^ with CS^D^ revealed 26 and 13 novel *Fatima* copies in CS^D^ and *Aet*^*D*^. These values were also quite similar when comparing CS with tetraploids (Fig. 4). For instance, *T. dicoccoides*^*AB*^ carries 474 (on A) and 478 (on B) novel copies of RLG_famc15 (*Jeli*) absent at orthologous loci CS^AB^, but, reversely, CS^AB^ carries 571 and 511 novel *Jeli* copies that are absent at orthologous loci in *T. dicoccoides*^*AB*^. Such unexpected similarity is not limited to *Fatima* and *Jeli* but rather true for almost all the other families in all species. However, we found a case, RLG_famc3 (*Laura*), that deviates from this pattern since it transposed 10 times more in CS^A^ than in *T. urartu*^*A*^ (1202 versus 113 specific insertions, respectively). This makes us conclude that, if the global transposition rate appears constant during the recent *Triticeae* history, each family amplified at a specific rate, which is not simply explained by its abundance, but this rate tends to remain constant across speciation events. Polyploidy is not associated with more copies accumulated over time and transposition rate of each family is not even disturbed following polyploidization events i.e., diploids, tetraploids, and hexaploids accumulated novel TEs at similar rates. Globally, all TE families generate novel copies, independently, at different genomic locations, but at approximately the same rate, in the different lineages.

**Figure 4:**
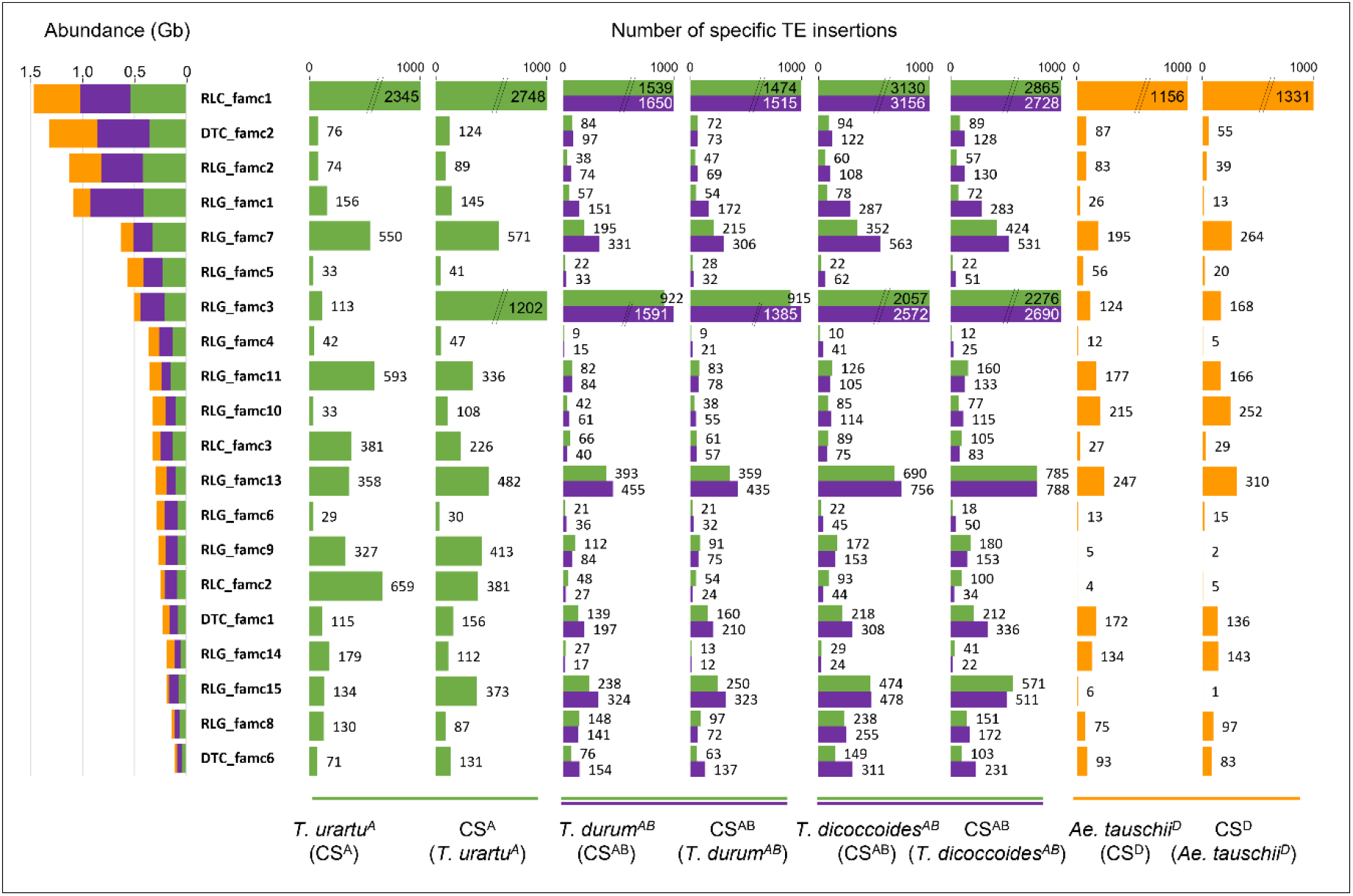
Histograms of the number of recent TE insertions discovered by whole genome alignments for the 20 most abundant wheat TE families. Abundance (in Gb) of TE families annotated in CS A, B, and D subgenomes (IWGSC RefSeq v1.1) is represented on the left panel. Specific TE insertions were identified in pairwise oIGR alignments and the 8 histograms represent the number of TEs that are present the query genome and absent at the orthologous locus in the compared genome which is mentioned in parentheses. Numbers of insertions are provided by subgenome with the following color code: A: green; B: violet; D: orange.

### Proximity to genes and inconsistent estimates of insertion dates

In total, our bp-resolved alignments revealed the presence of 51,928 elements (39,966 class 1, 11,048 class 2, and 914 unclassified) in the Chinese Spring genome that are absent from the corresponding insertion site in one or more compared genomes. They represent a large and clean dataset of novel insertions, although some of the class 2 elements detected here may be traces of recent excision in the compared genome. We used this dataset in order to investigate the global distribution of new insertions and the potential preferential insertion in gene vicinity. Among the 102,601 analyzed oIGRs, 26,862 contained one or more new insertions, showing that there was no insertion hotspot but rather that novel insertions spread the genome homogeneously. In wheat, genes tend to be clustered into small islands separated by large TE clusters, so it was important to distinguish small IGRs (<=40 kb; 51,120 regions corresponding to gene islands) from large ones (>40 kb; 51,481 regions corresponding to large TE clusters). Small IGRs represent only 4% of the TE space but accumulated 10% of the novel insertions: 5414 recent insertions scattered into 4530 small IGRs. There were also 46,514 novel insertions in 22,332 distinct large IGRs with, again, a number of novel insertions in the close vicinity of genes higher than expected by random insertion: 11% of insertions occurred within 5 kb around genes, i.e., 4% of target sequences. Affinity with genic regions depends on the TE family considered. We thus analyzed the insertional behavior for 44 TE families for which we had at least 100 novel insertions detected in CS. Half of them exhibit preferential insertion around genes (at least 3 times more insertions close to genes than expected randomly). They belong to class 2 transposons and LINEs retrotransposon families that were previously shown to be enriched in gene promoters (Wicker et al., 2018). In contrast, Gypsy and Copia insertions tend to be excluded from the gene vicinity and insert preferentially in the core of large IGRs.

Finally, we estimated the age of these novel insertions with the approach that is widely used by the community: aligning 5’ and 3’ Long Terminal Repeats (LTRs) of LTR-retrotransposons (LTR-RTs) and applying a molecular clock, considering that both LTRs are replicated (and thus identical) during transposition process. The purpose was to validate the reliability of this approach since we have sampled here the largest dataset of recently inserted LTR-RTs. The average insertion time estimated for the 88,269 newly inserted LTR-RTs identified in all our comparisons, is 590,000 years. Only 4% are estimated to be younger than 100,000 years ago. This estimate is surprisingly older than expected and not in accordance with the divergence time estimated for these species/accessions. Thus, we selected 209 LTR-RTs that are strictly specific to CS while absent from all other species/accessions, to ensure we collected very recent ones (hundreds/thousands of years). Actually, only 5/209 carry identical LTRs while all others already exhibit sequence differences. Moreover, 95% (198/209) are estimated to be older than 100,000 years, an inconsistency with the fact that they transposed specifically in CS. This raises serious doubts about the error-less replication of LTRs during insertion process in *Triticeae* which may have implications for our understanding of TE evolution.

## Discussion

Assembling the wheat genome has long been a challenge but we now have reached the pangenomics area with multiple high-quality genome assemblies available. SNP diversity was intensively characterized in order to get a world-wide view of the *Triticum* population structure, impact of selection, introgressions (Balfourier et al., 2019; Zhou et al., 2020), and to even build haplotype maps for genotype imputations (Brinton et al., 2020; Jordan et al., 2022). SNPs are easy to discover, to genotype, because bioinformatics pipelines that handle short-reads are well established and because technology advances tackle the complexity of the genome. However, lots remain to be done to go beyond the type of diversity we could investigate with SNPs which is basically no more than allele combinations. This is the goal of pangenomics which relies on discovering the hidden diversity with loci under presence/absence variations. For that, it is important to go beyond the “uniquely mappable area” of short-read based bioinformatics, especially when studying complex genomes. Structural variations (SVs) were only poorly characterized for wheat genes and partially addressed for TEs (Montenegro et al., 2017; De Oliveira et al., 2020). SVs are, however, of major importance to understand the molecular factors responsible for phenotypic variations and to understand evolutionary rules governing TE dynamics. Here, we faced the challenge of characterizing variability affecting Gbs of repeated elements in one of the most complex genomes ever assembled. Addressing this question required dedicated tools, strategies, and expertise in TE classification and annotation. We homogenized TE annotations of all available genome sequences using CLARITE and ClariTeRep library that were specifically developed for modeling wheat TEs (Daron et al., 2014) and previously used to comparing A-B-D TE content (Wicker et al., 2018). TE annotations are actually very dependent on the tools and library used so that it is hazardous to compare annotations performed by different groups. Most striking difference comes from CACTA transposons because the ClariTeRep library is enriched in CACTAs manually curated (Choulet et al., 2010) that are generally absent from other wheat specific TE libraries. Indeed, several *Triticeae* sequencing projects concluded that CACTAs represent 5-6% of the genome (Jia et al., 2013; Ling et al., 2013), while their proportion is around 15% when using ClariTeRep.

We established a workflow in order to compute base pair-resolved whole-genome sequence alignments for Gb-sized genomes, by taking advantage of the high level of gene collinearity of *Triticeae* genomes to anchor the alignments. We split the subgenomes into ∼30,000 intervals corresponding to individual orthologous intergenic regions flanked by pairs of collinear orthologs, thus reducing the alignment space so that most TEs are not repeated within the aligned interval. This, however, excluded noncollinear regions from the analyses and it was important to check whether such a filter introduces bias by excluding regions that may be more variable than the genome average. This is why we checked with unbiased estimators of variability: comparing genome-wide family proportions and mapping of all ISBP markers. Results from oIGR alignments were consistent with these unbiased estimates. Firstly, TE families showing copy number differences at the whole genome level were also the ones enriched in the variable TE fraction defined by the alignments. Secondly, the extent of variability defined by oIGR alignments were fully in line with the unbiased estimates based on ISBP mapping although slightly (2% on average) lower, confirming that noncollinear (non-aligned) regions contain more SVs than the genome average. However, this quality control demonstrated that we can be confident that the conclusions reached by interval-based whole genome alignments are sound.

Our conclusions are in favor of an equilibrium model and do not fit the model of TE evolution by cycles of bursts/silencing. First pieces of evidence that wheat TE dynamics do not follow this model was brought by comparing A with B and D subgenomes (Wicker et al., 2018). These lineages diverged millions of years ago so that TE turnover is nearly complete i.e., there are (almost) no more orthologous TEs at that scale. Although TEs evolved independently in diploids, many striking features appeared: the wide majority of TE families maintained their ancestral copy-number with abundant families remaining abundant, and low-copy families remaining at low-copy number in the three lineages. Moreover, families that tend to be enriched close to genes kept this behavior in all lineages, although novel copies did not target the same genes. All features led to conclude that TE dynamics was highly regulated and suggested evolutionary constraints to maintain global equilibrium. This is this model that we temptingly challenged here by characterizing the initial events of the ongoing TE turnover in three *Triticeae* lineages and the impact of polyploidy on TE activity. Our data revealed that the TE composition of the variable TE fraction resembles the ancestral conserved one. This confirms that TE turnover does not modify the global TE landscape. In contrast, it occurs while maintaining an equilibrium with unchanged TE family proportions. We did not see any burst of a given family nor families that would have decreased in proportion because of being completely silenced.

Rare cases of families that contribute more than others to recent genome diversification were observed. Interestingly, this was the case for subtelomeric satellite repeats (XXX_famc10) and centromeric retrotransposons Cereba/Quinta (RLG_famc8). This suggests that such elements that play a major structural role in shaping chromosome architecture comprise the most rapidly evolving elements of the genome.

Specific regions we identified originate from both TE deletions and insertions. An important outcome is that, whatever the genomes compared, whatever the ploidy levels, the proportion of specific sequences was similar in both the query and reference. We did not see a species that would have accumulated more novel insertions than others. The TE turnover seems to occur at a conserved rate in the different lineage which is, again, in favor of the equilibrium model with a controlled genome size. This strongly suggests that TE deletions compensate genome expansion by novel insertions over time. Since we reached a base pair-resolution by BLAST alignments, we were able to search for SVs that are traces of a recent insertion/excision: a complete TE with TSDs in the query genome *versus* an empty insertion site in the reference genome. Although we found thousands of such insertions, their accumulated size reached only 10% of the specific fraction (in bps), while we would expect close to 50%. The reason for this is most probably because criteria to call a novel insertion were too strict to avoid false positives. In addition, subsequent events of deletions/rearrangements either in the query or reference may have erased molecular traces of such recent insertions.

Another interesting result is that polyploids have not experienced more transpositions than their diploid relatives. Diploids and polyploids accumulated a very similar number of TE insertions per subgenome, confirming there was no genomic shock that would have been followed by TE deregulation. In contrast, our results tend to show that the rate of TE transposition and, thus, TE turnover, is stable over time and whole genome duplications did not destabilize this equilibrium. However, this model is not in agreement with the conclusions of a differential accumulation of TE families in *Triticum/Aegilops* (Yaakov et al., 2013b; Keidar-Friedman et al., 2018). One of the main sources of such discrepancies is the definition of what we call a family. Previous studies observed lineage-specific accumulations of families whereas we observed an equilibrium. In fact, we agree that families evolved into variants i.e., subfamilies, that have accumulated independently in different lineages. But many families that were called by different names are actually related, sometimes can even be considered members of the same families. Deciphering these phylogenetic relations between copies is crucial to understand that, even if it is possible to see variants overrepresented in one lineage, the important and striking result is that its family remained at an equilibrium in all lineages. The importance of connecting elements was also underlined in Stritt et al. (2021) where it has been shown that a rare TE family tend to maintained vertically at low-copy number throughout angiosperms, suggesting an alternative scenario prevailing the usual view of TE dynamics. Also similar to what we found, TE insertion polymorphisms explored in 53 *Brachypodium* accessions did not reveal any lineage-specific TE activations but rather a homogeneous activity and a stable transposition rate among populations, suggesting a conserved regulatory mechanism (Stritt et al., 2018). TE proliferation in *Triticeae* does not appear to be the result of an induction by the environment but rather a highly regulated process playing a role in basic genome functioning. This challenges the previous view of TEs as “invasive” elements as it becomes more and more suggested in literature by the perspective of the genome ecology where TEs would persist by creating their own niche (Kremer et al., 2020). This has been recently commented in maize where the analysis of family-level ecology of the genome led to conclude that, like in our study, most families have been active recently, each family having its own survival strategy representing the evolution of a distinct ecological niche (Stitzer et al., 2021). Despite the short evolutionary time frame studied here, we discovered traces of novel insertions for 346 families i.e., meaning that virtually all families are active actually. Same conclusions were reached in rice where ca. 11,000 new insertions of 251 families were detected in genomes of 3000 natural rice accessions (Liu et al., 2021). This highlighted the discrepancies between artificial stress-induced transposition and activity under natural conditions which provide two different views. Similarly, we wondered why our results are not in agreement with TE behavior observed in synthetic newly formed polyploid wheats where some TEs were mobilized in response to polyploidy (Yaakov and Kashkush, 2012; Yaakov et al., 2013b). As suggested earlier, instability following polyploidy is unlikely to be selected for under natural conditions (Zhao et al., 2011) and we support the hypothesis that only polyploids for which the equilibrium remained stable maintained across the generations.

Finally, the fact that recently inserted LTR-RTs did not carry identical LTRs made us underline some doubts about the method used to estimate dates of insertions. Similar inconsistencies were reported based on the analysis of ca. 25,000 LTR-RTs in fifteen plant species which revealed that one fourth of nested retrotransposons exhibited older insertion date than pre-existing elements in which they had been inserted (Jedlicka et al., 2020). Gene conversion was suggested to be involved in erasing divergence over time. Another possible explanation would be that replication of LTRs during the insertion mechanism is error-prone. This may have strong impact on the way the community uses the molecular clock to date insertion events.

## Supplementary information

Supplemental Table S1: Proportions of TE superfamilies in the A, B, and D subgenomes of the *T. aestivum* genomes annotated with CLARITE.

## Availability of supporting data

Data generated (GFFs files of TE annotation, BED files of positions of orthologous regions aligned, positions of variable regions, and positions of novel insertions) were deposited in https://entrepot.recherche.data.gouv.fr under the doi:10.57745/RCTOQM.

Scripts used are available on Gitlab at https://forgemia.inra.fr/umr-gdec/scripts_files/

## Acknowledgments

Computations have been performed on the supercomputer facilities of the Mésocentre Clermont Auvergne University.

## Author Approvals

All authors have seen and approved the manuscript. Manuscript has not been accepted or published elsewhere.

## Competing interests

The authors declare that they have no competing interests.

## Funding

NP PhD was financed by a grant from the French Ministry of Higher Education, Research and Innovation (MESRI).

## Author’s contributions

NP downloaded the data material and performed most of the bioinformatic analyses. PL performed the analysis of TE insertions relatively to the closest gene. HR and RDO performed bioinformatic analyses related to genome annotations and file formatting. FC performed TE family enrichment analysis. FC, NP, and PL wrote the manuscript. EP and FC supervised the study.

## Notes

### Competing Interest Statement

The authors have declared no competing interest.

## References

Aury, J.M., Engelen, S., Istace, B., Monat, C., Lasserre-Zuber, P., Belser, C., Cruaud, C., Rimbert, H., Leroy, P., Arribat, S., Dufau, I., Bellec, A., Grimbichler, D., Papon, N., Paux, E., Ranoux, M., Alberti, A., Wincker, P., and Choulet, F. (2022). Long-read and chromosome-scale assembly of the hexaploid wheat genome achieves high resolution for research and breeding. Gigascience 11.

Avni, R., Lux, T., Minz-Dub, A., Millet, E., Sela, H., Distelfeld, A., Deek, J., Yu, G., Steuernagel, B., Pozniak, C., Ens, J., Gundlach, H., Mayer, K.F.X., Himmelbach, A., Stein, N., Mascher, M., Spannagl, M., Wulff, B.B.H., and Sharon, A. (2022). Genome sequences of three Aegilops species of the section Sitopsis reveal phylogenetic relationships and provide resources for wheat improvement. Plant J 110, 179–192.

Avni, R., Nave, M., Barad, O., Baruch, K., Twardziok, S.O., Gundlach, H., Hale, I., Mascher, M., Spannagl, M., Wiebe, K., Jordan, K.W., Golan, G., Deek, J., Ben-Zvi, B., Ben-Zvi, G., Himmelbach, A., MacLachlan, R.P., Sharpe, A.G., Fritz, A., Ben-David, R., Budak, H., Fahima, T., Korol, A., Faris, J.D., Hernandez, A., Mikel, M.A., Levy, A.A., Steffenson, B., Maccaferri, M., Tuberosa, R., Cattivelli, L., Faccioli, P., Ceriotti, A., Kashkush, K., Pourkheirandish, M., Komatsuda, T., Eilam, T., Sela, H., Sharon, A., Ohad, N., Chamovitz, D.A., Mayer, K.F.X., Stein, N., Ronen, G., Peleg, Z., Pozniak, C.J., Akhunov, E.D., and Distelfeld, A. (2017). Wild emmer genome architecture and diversity elucidate wheat evolution and domestication. Science 357, 93–97.

Baduel, P., and Quadrana, L. (2021). Jumpstarting evolution: How transposition can facilitate adaptation to rapid environmental changes. Curr Opin Plant Biol 61, 102043.

Balfourier, F., Bouchet, S., Robert, S., De Oliveira, R., Rimbert, H., Kitt, J., Choulet, F., Paux, E., Consortium, I.W.G.S., and Consortium, B. (2019). Worldwide phylogeography and history of wheat genetic diversity. Sci Adv 5, eaav0536.

Bariah, I., Keidar-Friedman, D., and Kashkush, K. (2020). Where the Wild Things Are: Transposable Elements as Drivers of Structural and Functional Variations in the Wheat Genome. Front Plant Sci 11, 585515.

Brinton, J., Ramirez-Gonzalez, R.H., Simmonds, J., Wingen, L., Orford, S., Griffiths, S., Haberer, G., Spannagl, M., Walkowiak, S., Pozniak, C., Uauy, C., and Project, W.G. (2020). A haplotype-led approach to increase the precision of wheat breeding. Commun Biol 3, 712.

Camacho, C., Coulouris, G., Avagyan, V., Ma, N., Papadopoulos, J., Bealer, K., and Madden, T.L. (2009). BLAST+: architecture and applications. BMC Bioinformatics 10, 421.

Choulet, F., Wicker, T., Rustenholz, C., Paux, E., Salse, J., Leroy, P., Schlub, S., Le Paslier, M.C., Magdelenat, G., Gonthier, C., Couloux, A., Budak, H., Breen, J., Pumphrey, M., Liu, S., Kong, X., Jia, J., Gut, M., Brunel, D., Anderson, J.A., Gill, B.S., Appels, R., Keller, B., and Feuillet, C. (2010). Megabase level sequencing reveals contrasted organization and evolution patterns of the wheat gene and transposable element spaces. Plant Cell 22, 1686–1701.

Cosby, R.L., Chang, N.C., and Feschotte, C. (2019). Host-transposon interactions: conflict, cooperation, and cooption. Genes Dev 33, 1098–1116.

Daron, J., Glover, N., Pingault, L., Theil, S., Jamilloux, V., Paux, E., Barbe, V., Mangenot, S., Alberti, A., Wincker, P., Quesneville, H., Feuillet, C., and Choulet, F. (2014). Organization and evolution of transposable elements along the bread wheat chromosome 3B. Genome Biol 15, 546.

De Oliveira, R., Rimbert, H., Balfourier, F., Kitt, J., Dynomant, E., Vrána, J., DoleŽel, J., Cattonaro, F., Paux, E., and Choulet, F. (2020). Structural Variations Affecting Genes and Transposable Elements of Chromosome 3B in Wheats. Front Genet 11, 891.

Dvorak, J., Wang, L., Zhu, T., Jorgensen, C.M., Luo, M.C., Deal, K.R., Gu, Y.Q., Gill, B.S., Distelfeld, A., Devos, K.M., Qi, P., and McGuire, P.E. (2018). Reassessment of the evolution of wheat chromosomes 4A, 5A, and 7B. Theor Appl Genet 131, 2451–2462.

Glémin, S., Scornavacca, C., Dainat, J., Burgarella, C., Viader, V., Ardisson, M., Sarah, G., Santoni, S., David, J., and Ranwez, V. (2019). Pervasive hybridizations in the history of wheat relatives. Sci Adv 5, eaav9188.

Guo, W., Xin, M., Wang, Z., Yao, Y., Hu, Z., Song, W., Yu, K., Chen, Y., Wang, X., Guan, P., Appels, R., Peng, H., Ni, Z., and Sun, Q. (2020). Origin and adaptation to high altitude of Tibetan semi-wild wheat. Nat Commun 11, 5085.

IWGSC. (2018). Shifting the limits in wheat research and breeding using a fully annotated reference genome. Science 361.

Jedlicka, P., Lexa, M., and Kejnovsky, E. (2020). What Can Long Terminal Repeats Tell Us About the Age of LTR Retrotransposons, Gene Conversion and Ectopic Recombination? Front Plant Sci 11, 644.

Jia, J., Zhao, S., Kong, X., Li, Y., Zhao, G., He, W., Appels, R., Pfeifer, M., Tao, Y., Zhang, X., Jing, R., Zhang, C., Ma, Y., Gao, L., Gao, C., Spannagl, M., Mayer, K.F., Li, D., Pan, S., Zheng, F., Hu, Q., Xia, X., Li, J., Liang, Q., Chen, J., Wicker, T., Gou, C., Kuang, H., He, G., Luo, Y., Keller, B., Xia, Q., Lu, P., Wang, J., Zou, H., Zhang, R., Xu, J., Gao, J., Middleton, C., Quan, Z., Liu, G., Yang, H., Liu, X., He, Z., Mao, L., and Consortium, I.W.G.S. (2013). Aegilops tauschii draft genome sequence reveals a gene repertoire for wheat adaptation. Nature 496, 91–95.

Jordan, K.W., Bradbury, P.J., Miller, Z.R., Nyine, M., He, F., Fraser, M., Anderson, J., Mason, E., Katz, A., Pearce, S., Carter, A.H., Prather, S., Pumphrey, M., Chen, J., Cook, J., Liu, S., Rudd, J.C., Wang, Z., Chu, C., Ibrahim, A.M.H., Turkus, J., Olson, E., Nagarajan, R., Carver, B., Yan, L., Taagen, E., Sorrells, M., Ward, B., Ren, J., Akhunova, A., Bai, G., Bowden, R., Fiedler, J., Faris, J., Dubcovsky, J., Guttieri, M., Brown-Guedira, G., Buckler, E., Jannink, J.L., and Akhunov, E.D. (2022). Development of the Wheat Practical Haplotype Graph database as a resource for genotyping data storage and genotype imputation. G3 (Bethesda) 12.

Keidar-Friedman, D., Bariah, I., and Kashkush, K. (2018). Genome-wide analyses of miniature inverted-repeat transposable elements reveals new insights into the evolution of the Triticum-Aegilops group. PLoS One 13, e0204972.

Kraitshtein, Z., Yaakov, B., Khasdan, V., and Kashkush, K. (2010). Genetic and epigenetic dynamics of a retrotransposon after allopolyploidization of wheat. Genetics 186, 801–812.

Kremer, S.C., Linquist, S., Saylor, B., Elliott, T.A., Gregory, T.R., and Cottenie, K. (2020). Transposable element persistence via potential genome-level ecosystem engineering. BMC Genomics 21, 367.

Levy, A.A., and Feldman, M. (2022). Evolution and origin of bread wheat. Plant Cell.

Li, H. (2018). Minimap2: pairwise alignment for nucleotide sequences. Bioinformatics 34, 3094–3100.

Li, L.F., Zhang, Z.B., Wang, Z.H., Li, N., Sha, Y., Wang, X.F., Ding, N., Li, Y., Zhao, J., Wu, Y., Gong, L., Mafessoni, F., Levy, A.A., and Liu, B. (2022). Genome sequences of five Sitopsis species of Aegilops and the origin of polyploid wheat B subgenome. Mol Plant 15, 488–503.

Ling, H.Q., Ma, B., Shi, X., Liu, H., Dong, L., Sun, H., Cao, Y., Gao, Q., Zheng, S., Li, Y., Yu, Y., Du, H., Qi, M., Lu, H., Yu, H., Cui, Y., Wang, N., Chen, C., Wu, H., Zhao, Y., Zhang, J., Zhou, W., Zhang, B., Hu, W., van Eijk, M.J.T., Tang, J., Witsenboer, H.M.A., Zhao, S., Li, Z., Zhang, A., Wang, D., and Liang, C. (2018). Genome sequence of the progenitor of wheat A subgenome Triticum urartu. Nature 557, 424–428.

Ling, H.Q., Zhao, S., Liu, D., Wang, J., Sun, H., Zhang, C., Fan, H., Li, D., Dong, L., Tao, Y., Gao, C., Wu, H., Li, Y., Cui, Y., Guo, X., Zheng, S., Wang, B., Yu, K., Liang, Q., Yang, W., Lou, X., Chen, J., Feng, M., Jian, J., Zhang, X., Luo, G., Jiang, Y., Liu, J., Wang, Z., Sha, Y., Zhang, B., Tang, D., Shen, Q., Xue, P., Zou, S., Wang, X., Liu, X., Wang, F., Yang, Y., An, X., Dong, Z., Zhang, K., Luo, M.C., Dvorak, J., Tong, Y., Yang, H., Li, Z., Wang, D., and Zhang, A. (2013). Draft genome of the wheat A-genome progenitor Triticum urartu. Nature 496, 87–90.

Lisch, D. (2013). How important are transposons for plant evolution? Nat Rev Genet 14, 49–61.

Liu, Z., Zhao, H., Yan, Y., Wei, M.X., Zheng, Y.C., Yue, E.K., Alam, M.S., Smartt, K.O., Duan, M.H., and Xu, J.H. (2021). Extensively Current Activity of Transposable Elements in Natural Rice Accessions Revealed by Singleton Insertions. Front Plant Sci 12, 745526.

Luo, M.C., Gu, Y.Q., Puiu, D., Wang, H., Twardziok, S.O., Deal, K.R., Huo, N., Zhu, T., Wang, L., Wang, Y., McGuire, P.E., Liu, S., Long, H., Ramasamy, R.K., Rodriguez, J.C., Van, S.L., Yuan, L., Wang, Z., Xia, Z., Xiao, L., Anderson, O.D., Ouyang, S., Liang, Y., Zimin, A.V., Pertea, G., Qi, P., Bennetzen, J.L., Dai, X., Dawson, M.W., Müller, H.G., Kugler, K., Rivarola-Duarte, L., Spannagl, M., Mayer, K.F.X., Lu, F.H., Bevan, M.W., Leroy, P., Li, P., You, F.M., Sun, Q., Liu, Z., Lyons, E., Wicker, T., Salzberg, S.L., Devos, K.M., and Dvořák, J. (2017). Genome sequence of the progenitor of the wheat D genome Aegilops tauschii. Nature 551, 498–502.

Maccaferri, M., Harris, N.S., Twardziok, S.O., Pasam, R.K., Gundlach, H., Spannagl, M., Ormanbekova, D., Lux, T., Prade, V.M., Milner, S.G., Himmelbach, A., Mascher, M., Bagnaresi, P., Faccioli, P., Cozzi, P., Lauria, M., Lazzari, B., Stella, A., Manconi, A., Gnocchi, M., Moscatelli, M., Avni, R., Deek, J., Biyiklioglu, S., Frascaroli, E., Corneti, S., Salvi, S., Sonnante, G., Desiderio, F., Marè, C., Crosatti, C., Mica, E., Özkan, H., Kilian, B., De Vita, P., Marone, D., Joukhadar, R., Mazzucotelli, E., Nigro, D., Gadaleta, A., Chao, S., Faris, J.D., Melo, A.T.O., Pumphrey, M., Pecchioni, N., Milanesi, L., Wiebe, K., Ens, J., MacLachlan, R.P., Clarke, J.M., Sharpe, A.G., Koh, C.S., Liang, K.Y.H., Taylor, G.J., Knox, R., Budak, H., Mastrangelo, A.M., Xu, S.S., Stein, N., Hale, I., Distelfeld, A., Hayden, M.J., Tuberosa, R., Walkowiak, S., Mayer, K.F.X., Ceriotti, A., Pozniak, C.J., and Cattivelli, L. (2019). Durum wheat genome highlights past domestication signatures and future improvement targets. Nat Genet 51, 885–895.

Marcussen, T., Sandve, S.R., Heier, L., Spannagl, M., Pfeifer, M., International Wheat Genome Sequencing, C., Jakobsen, K.S., Wulff, B.B., Steuernagel, B., Mayer, K.F., and Olsen, O.A. (2014). Ancient hybridizations among the ancestral genomes of bread wheat. Science 345, 1250092.

Middleton, C.P., Senerchia, N., Stein, N., Akhunov, E.D., Keller, B., Wicker, T., and Kilian, B. (2014). Sequencing of chloroplast genomes from wheat, barley, rye and their relatives provides a detailed insight into the evolution of the Triticeae tribe. PLoS One 9, e85761.

Montenegro, J.D., Golicz, A.A., Bayer, P.E., Hurgobin, B., Lee, H., Chan, C.K., Visendi, P., Lai, K., DoleŽel, J., Batley, J., and Edwards, D. (2017). The pangenome of hexaploid bread wheat. Plant J 90, 1007–1013.

Paux, E., Roger, D., Badaeva, E., Gay, G., Bernard, M., Sourdille, P., and Feuillet, C. (2006). Characterizing the composition and evolution of homoeologous genomes in hexaploid wheat through BAC-end sequencing on chromosome 3B. Plant J 48, 463–474.

Quinlan, A.R., and Hall, I.M. (2010). BEDTools: a flexible suite of utilities for comparing genomic features. Bioinformatics 26, 841–842.

SanMiguel, P., Gaut, B.S., Tikhonov, A., Nakajima, Y., and Bennetzen, J.L. (1998). The paleontology of intergene retrotransposons of maize. Nat Genet 20, 43–45.

Smit, A.F.A., Hubley, R., and Green, P. (1996-2004). RepeatMasker Open-3.0 http://www.repeatmasker.org.

Stitzer, M.C., Anderson, S.N., Springer, N.M., and Ross-Ibarra, J. (2021). The genomic ecosystem of transposable elements in maize. PLoS Genet 17, e1009768.

Stritt, C., Thieme, M., and Roulin, A.C. (2021). Rare transposable elements challenge the prevailing view of transposition dynamics in plants. Am J Bot 108, 1310–1314.

Stritt, C., Gordon, S.P., Wicker, T., Vogel, J.P., and Roulin, A.C. (2018). Recent Activity in Expanding Populations and Purifying Selection Have Shaped Transposable Element Landscapes across Natural Accessions of the Mediterranean Grass Brachypodium distachyon. Genome Biol Evol 10, 304–318.

Walkowiak, S., Gao, L., Monat, C., Haberer, G., Kassa, M.T., Brinton, J., Ramirez-Gonzalez, R.H., Kolodziej, M.C., Delorean, E., Thambugala, D., Klymiuk, V., Byrns, B., Gundlach, H., Bandi, V., Siri, J.N., Nilsen, K., Aquino, C., Himmelbach, A., Copetti, D., Ban, T., Venturini, L., Bevan, M., Clavijo, B., Koo, D.H., Ens, J., Wiebe, K., N’Diaye, A., Fritz, A.K., Gutwin, C., Fiebig, A., Fosker, C., Fu, B.X., Accinelli, G.G., Gardner, K.A., Fradgley, N., Gutierrez-Gonzalez, J., Halstead-Nussloch, G., Hatakeyama, M., Koh, C.S., Deek, J., Costamagna, A.C., Fobert, P., Heavens, D., Kanamori, H., Kawaura, K., Kobayashi, F., Krasileva, K., Kuo, T., McKenzie, N., Murata, K., Nabeka, Y., Paape, T., Padmarasu, S., Percival-Alwyn, L., Kagale, S., Scholz, U., Sese, J., Juliana, P., Singh, R., Shimizu-Inatsugi, R., Swarbreck, D., Cockram, J., Budak, H., Tameshige, T., Tanaka, T., Tsuji, H., Wright, J., Wu, J., Steuernagel, B., Small, I., Cloutier, S., Keeble-Gagnère, G., Muehlbauer, G., Tibbets, J., Nasuda, S., Melonek, J., Hucl, P.J., Sharpe, A.G., Clark, M., Legg, E., Bharti, A., Langridge, P., Hall, A., Uauy, C., Mascher, M., Krattinger, S.G., Handa, H., Shimizu, K.K., Distelfeld, A., Chalmers, K., Keller, B., Mayer, K.F.X., Poland, J., Stein, N., McCartney, C.A., Spannagl, M., Wicker, T., and Pozniak, C.J. (2020). Multiple wheat genomes reveal global variation in modern breeding. Nature 588, 277–283.

Wicker, T., Stritt, C., Sotiropoulos, A.G., Poretti, M., Pozniak, C., Walkowiak, S., Gundlach, H., and Stein, N. (2022). Transposable Element Populations Shed Light on the Evolutionary History of Wheat and the Complex Co-Evolution of Autonomous and Non-Autonomous Retrotransposons. Advanced Genetics 3, 2100022.

Wicker, T., Gundlach, H., Spannagl, M., Uauy, C., Borrill, P., Ramirez-Gonzalez, R.H., De Oliveira, R., International Wheat Genome Sequencing, C., Mayer, K.F.X., Paux, E., and Choulet, F. (2018). Impact of transposable elements on genome structure and evolution in bread wheat. Genome Biol 19, 103.

Wicker, T., Sabot, F., Hua-Van, A., Bennetzen, J.L., Capy, P., Chalhoub, B., Flavell, A., Leroy, P., Morgante, M., Panaud, O., Paux, E., SanMiguel, P., and Schulman, A.H. (2007). A unified classification system for eukaryotic transposable elements. Nat Rev Genet 8, 973–982.

Wu, T.D., and Watanabe, C.K. (2005). GMAP: a genomic mapping and alignment program for mRNA and EST sequences. Bioinformatics 21, 1859–1875.

Yaakov, B., and Kashkush, K. (2011). Massive alterations of the methylation patterns around DNA transposons in the first four generations of a newly formed wheat allohexaploid. Genome 54, 42–49.

Yaakov, B., and Kashkush, K. (2012). Mobilization of Stowaway-like MITEs in newly formed allohexaploid wheat species. Plant Mol Biol 80, 419–427.

Yaakov, B., Ben-David, S., and Kashkush, K. (2013a). Genome-wide analysis of Stowaway-like MITEs in wheat reveals high sequence conservation, gene association, and genomic diversification. Plant Physiol 161, 486–496.

Yaakov, B., Meyer, K., Ben-David, S., and Kashkush, K. (2013b). Copy number variation of transposable elements in Triticum-Aegilops genus suggests evolutionary and revolutionary dynamics following allopolyploidization. Plant Cell Rep 32, 1615–1624.

Zhao, N., Zhu, B., Li, M., Wang, L., Xu, L., Zhang, H., Zheng, S., Qi, B., Han, F., and Liu, B. (2011). Extensive and heritable epigenetic remodeling and genetic stability accompany allohexaploidization of wheat. Genetics 188, 499–510.

Zhou, Y., Zhao, X., Li, Y., Xu, J., Bi, A., Kang, L., Xu, D., Chen, H., Wang, Y., Wang, Y.G., Liu, S., Jiao, C., Lu, H., Wang, J., Yin, C., Jiao, Y., and Lu, F. (2020). Triticum population sequencing provides insights into wheat adaptation. Nat Genet 52, 1412–1422.

Zhu, T., Wang, L., Rodriguez, J.C., Deal, K.R., Avni, R., Distelfeld, A., McGuire, P.E., Dvorak, J., and Luo, M.C. (2019). Improved Genome Sequence of Wild Emmer Wheat Zavitan with the Aid of Optical Maps. G3 (Bethesda) 9, 619–624.

